# LactoTypeDB: a regenerable, type-anchored 16S rRNA gene reference for species-level identification of the *Lactobacillaceae* in foods

**DOI:** 10.64898/2026.08.12.744342

**Authors:** Scott A. Oliphant, Jennifer M. Gardner, Vladimir Jiranek, Krista M. Sumby

## Abstract

Amplicon surveys of fermented and spoiled foods routinely resolve *Lactobacillaceae*, the lactic acid bacteria responsible for many food and beverage fermentations, only to genus, whereas registers such as the Inventory of Microbial Food Cultures require species-level identification. This shortfall arises from the 16S rRNA gene’s limited, region-dependent resolution and from incomplete, non-type-strain-anchored references that silently reassign missing species to their nearest relative. We built LactoTypeDB, a regenerable, type-anchored reference covering 434 of the family’s 441 species and all 37 genera and substituted it into the Living Tree Project release LTP 08_2023 the field’s default classifier uses. This eliminated species-level misassignment of type strains in all regions tested and cut misassignment of 10,329 other sequences from the same species from 1,374 errors down to 3 when the full-length 16S rRNA gene was used. Applied unmodified to 11,612 V3-V4 distinct sequences from a published survey of two meat production lines, the workflow returned a species for 213 and a genus for 5,926, and flagged 3,495 as undescribed candidates, more than a third of them nearest to *Dellaglioa*, a genus that includes a meat-spoilage organism tracked in that survey. The ambiguity that remains is the marker’s, since V3-V4 collapses 417 of the 434 species into 27 groups it cannot separate. For food microbiology laboratories, the practical change is that a species call from this family can now be trusted where the marker allows it, and a sequence matching nothing becomes a candidate worth isolating rather than a limitation to work around.

## 1. INTRODUCTION

In 2020, Zheng et al. split the single genus *Lactobacillus* into 25 genera, 23 of them new, and emended the family *Lactobacillaceae* to include the genera of the *Leuconostocaceae*, a reclassification that outpaced the general-purpose reference databases used to identify its members (Zheng et al., 2020). These lactic acid bacteria ferment foods and beverages, preserve silage and are sold as probiotics. Some species also spoil chilled meat, where they dominate the community during storage (Poirier et al., 2023). The family is surveyed almost exclusively by 16S rRNA gene amplicon sequencing (Parente et al., 2023; Zheng et al., 2020). Such surveys usually stop at genus, a rank too coarse for regulatory and industrial use: species-rank identification is required for admission to lists such as the Inventory of Microbial Food Cultures, where this family accounts for 110 of 314 listed species (Parente et al., 2023). Importantly, the same rank decides safety status. The European Food Safety Authority grants qualified presumption of safety at species level for bacteria and re-verifies the names carrying that status against the nomenclatural authorities every six months. The most recent of those revisions moved *Lacticaseibacillus rhamnosus* to a synonym and made *Lactobacillus rhamnosus* the correct name (EFSA BIOHAZ Panel, 2026, which spells the genus *Lactibacillus* in each of the three places it states the change). That reverses the direction every other transfer in this family took, and it leaves one species under two names in current use, the one the 2020 reclassification gave it and the one LPSN now rules correct. The reference described here carries the LPSN name, by the rule of Section 2.1. A species-level misassignment therefore moves a strain across a regulatory boundary, and a survey reporting genus alone cannot show that it has happened.

Parente et al. (2023) tested how far such a survey can go, classifying the type strains’ own 16S rRNA gene sequences with the standard software (the dada2 package; Callahan et al., 2016; Callahan, 2025), against the SILVA reference database (Quast et al., 2013), at each method’s default settings. Genus was correctly assigned for 97.5% of full-length sequences, falling to 92.6% for the shorter V4 region. Species assignment was unambiguous for only 49.9% of full-length sequences and less for every shorter region, which Parente et al. (2023) attributed to the high 16S rRNA gene sequence similarity among closely related species of *Lactiplantibacillus* and *Lacticaseibacillus*.

Parente et al. (2023) established the scope of the problem this work addresses. By scoring the type strains against their own reference, they showed genus assignment performing well and species assignment performing poorly, and by reporting that shortfall region by region rather than as one overall figure, they made it possible to ask whether the problem lies with the marker or with the reference. A wider survey of the published food literature by Parente et al. (2022) reached the same conclusion from the opposite direction: genus can usually be assigned with confidence across the commonly used regions, but species-level calls need confirming. Neither of the reference databases that benchmark relied on is actively maintained, a limitation this study addresses. SILVA v138.1 carries no species-level labels for the genera that used to make up *Lactobacillus*, so Parente et al. (2023) built one, relabelling each SILVA sequence of the 23 new genera with the species of the type-strain sequence at smallest 5-mer distance. Those type-strain sequences were downloaded from LPSN in a single pass, and any sequence shorter than 1,300 base pairs, or carrying more than five ambiguous bases, was swapped for the equivalent record from GenBank or RefSeq. The result covered the 362 species formally described by the time of their publication. This generated a static list, applicable to that moment, rather than a dynamic one that can be regenerated as new species are described.

The family has since grown to 441 species, but the reference trees built to track all of bacterial taxonomy (Ludwig et al., 2021; Yarza et al., 2008) don’t track any single family to completion. The most widely used of these, the Living Tree Project (release LTP 08_2023), holds only 376 of the 434 species covered here. Its 2026 successor to that release, LTP+ 02_2026, is larger, holding 87,282 sequences drawn additionally from three other major databases (GTDB, SILVA and NCBI; Viver et al., 2026), but most of its records aren’t anchored to a type strain; only 21,376 are (Table S1). This matters because when a reference has no record for a query’s true species, standard classifiers don’t leave it unassigned: they silently reassign it to whichever represented species is the closest match, with nothing in the output flagging that a substitution happened, a failure mode inherent to classifying against an incomplete (“closed”) reference (Edgar, 2018a; Escapa et al., 2020; Rohwer et al., 2018). Using a stricter confidence threshold reduces how often this happens, but at the cost of leaving more queries unassigned altogether (Escapa et al., 2020).

No existing 16S rRNA gene reference for the *Lactobacillaceae* is both complete and able to be rebuilt as the family’s taxonomy keeps changing. This study set out to build one, to test whether doing so lets us tell apart two causes of ambiguity that had previously been tangled together, an incomplete reference versus a genuine limit of the marker itself, and to ask what a survey can still say about sequences that remain unnamed even with a complete reference in hand. We describe LactoTypeDB, a type-anchored reference in which every record derives from the type strain of the name it carries, and which one command rebuilds from a dated, verifiable snapshot of two nomenclature and sequence databases, LPSN and NCBI (Table 1, Table S2). We repeat the species-level measurement of Parente et al. (2023) and show why the two values are not directly comparable, and we extend their approach with a second classification step, run after the usual genus-level call: rule-out software from Tanes et al. (2024) that returns either the one species a sequence can’t be distinguished from, the set of species it’s compatible with, or nothing if it matches none of them. We then run that combined workflow, unmodified, using our deposited scripts, on 11,612 distinct sequences from a published survey of two fresh meat production lines (Poirier et al., 2023), presenting the method and its first real-world test together. This work contributes the reference database itself and the code that regenerates it. The workflow uses two established tools: the standard genus-assignment function from the dada2 package (Callahan et al., 2016) and the rule-out software of Tanes et al. (2024). Both are run at their published default settings, and neither is redistributed here. Beyond the list of names provided, nothing in the reference-building procedure is specific to this family, so the same approach can be applied to any bacterial family whose names and type-strain genomes are catalogued in LPSN and NCBI.

**TABLE 1.**
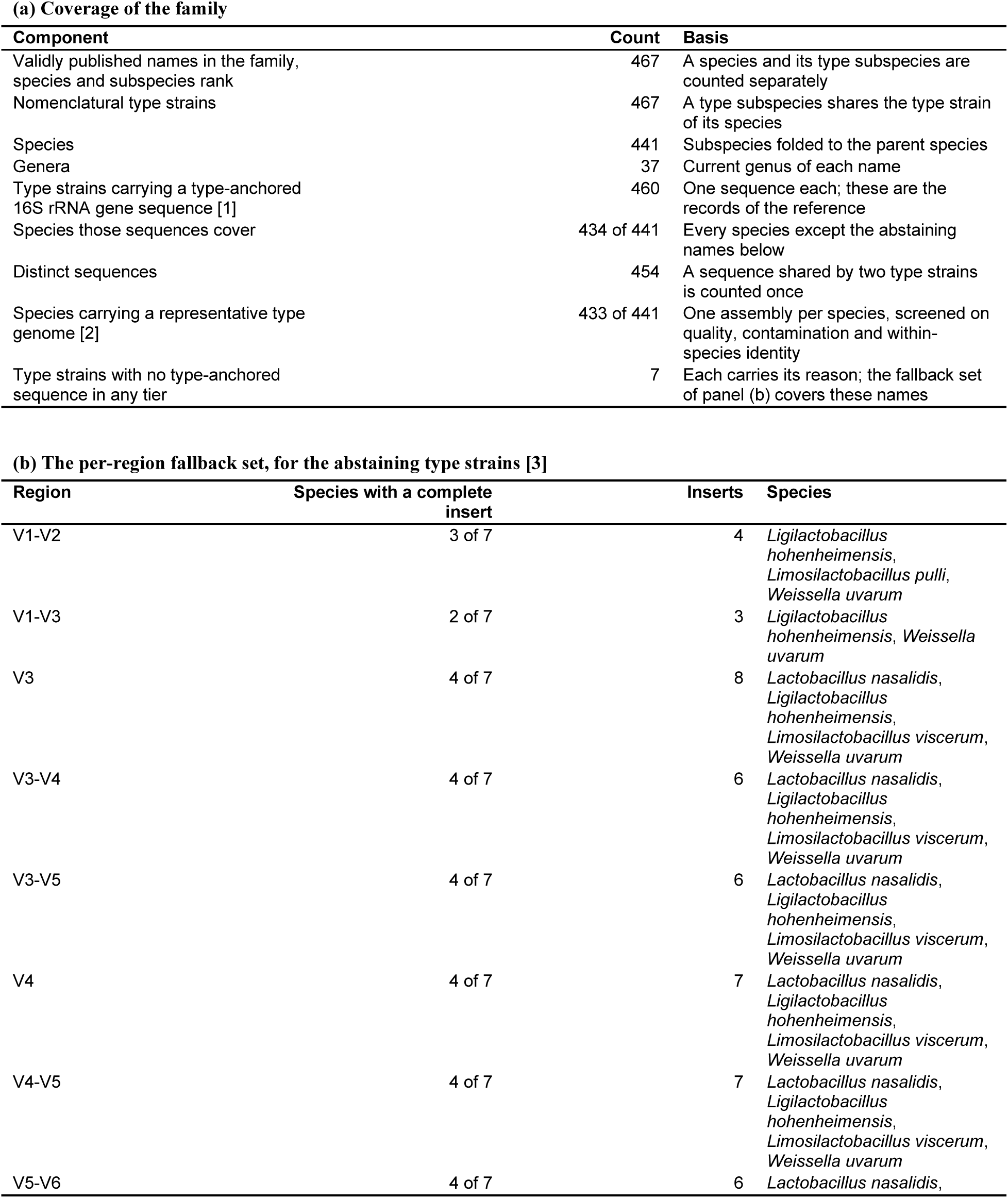
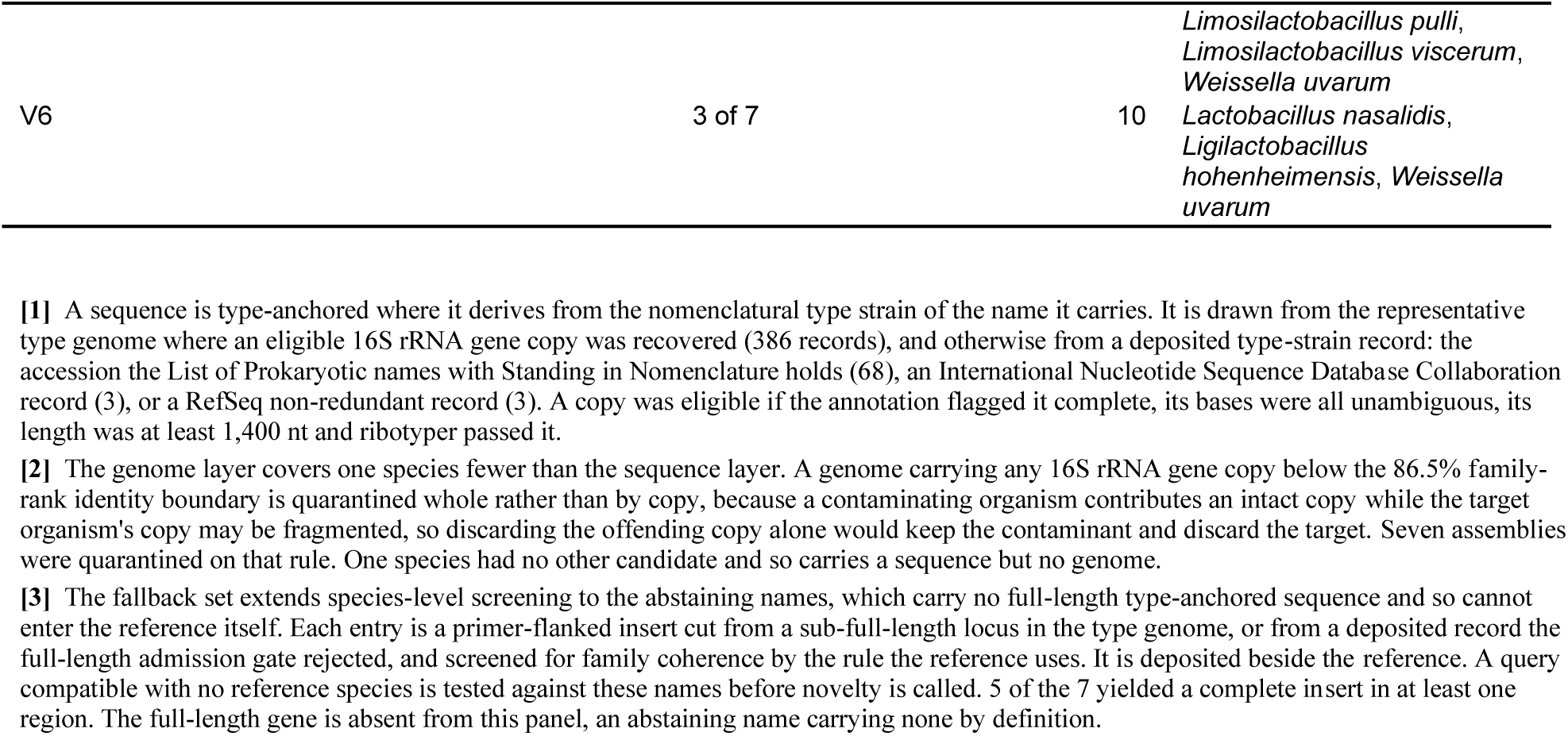
Coverage of LactoTypeDB v1.0.0, and the per-region fallback set for the names it cannot represent. Panel (a) gives the share of the family that carries a record, at each rank the question can be asked at. Panel (b) gives the fallback set. The seven abstaining type strains carry no full-length type-anchored sequence, so a per-region insert is deposited for them separately, and a query compatible with no reference species is tested against those names before novelty is called. Every count is measured from the deposited tables of release v1.0.0. The rule producing each is given in the third column of panel (a) and the qualifications in footnotes 1 to 3. The release, the code that builds it and the checksum-pinned snapshot of its source inputs are public, so each count is reproducible from the frozen input.

## 2. MATERIALS AND METHODS

### 2.1. Reference construction

Names were taken from LPSN by parent identifier, walking the *Lactobacillaceae* record (identifier 728) down to subspecies rank, rather than by name search, and matched against NCBI Taxonomy identifier 33958 on the identifier columns. Candidate genome assemblies designated as from type material were retrieved through NCBI Datasets (O’Leary et al., 2024) from the RefSeq and GenBank assembly summary reports, scoped to the NCBI Taxonomy subtree of identifier 33958 on the taxid columns, deduplicated on the numeric accession core, and restricted to those flagged as from type material, which gave the 829 candidates and their CheckM values and contig N50. An assembly was discarded where CheckM completeness fell below 90% or contamination exceeded 5%, so an assembly exactly on either boundary was kept, and one carrying no CheckM value was recorded as unknown rather than as a failure.

Every 16S rRNA gene copy of every admitted candidate was then screened for family coherence, before ranking rather than after it, and a candidate carrying any copy below the 86.5% family-rank identity floor was quarantined whole rather than by copy and dropped from selection (Table 1, footnote 2). A species whose top candidate is quarantined moves to its next best candidate.

Only a species with no remaining candidate is excluded from the genome layer. Retained candidates of each species were then ranked on five keys in order: a confirmed CheckM pass before an unknown, RefSeq before GenBank, complete genome before chromosome, scaffold and contig, higher contig N50, and lowest numeric accession core. The highest ranked was chosen as the representative. A 16S rRNA gene copy was eligible for inclusion if it was annotated as complete, carried no ambiguous bases, met a minimum length of 1,400 nt and passed quality control by ribotyper. Where a representative genome yielded no eligible copy, the sequence came from the deposited type-strain tiers of Table 1, footnote 1. One name is handled two ways, by design. LPSN rules *Lactobacillus rhamnosus* the correct name under a clinical-stability override and *Lacticaseibacillus rhamnosus* a suspended synonym, which is the opposite direction from every other transfer in this family. The deposited reference carries the LPSN name, so a survey run under this workflow reports *Lactobacillus rhamnosus*, which is the name EFSA’s most recent qualified presumption of safety revision also rules correct (Section 1). The roster used to score the benchmarks of Section 3.2 takes the phylogenetic name instead, because holding that species out and reclassifying its conspecifics placed 338 of 339 in *Lacticaseibacillus*. The difference affects 1 of the 434 species, is disclosed per record in Table S3, and is a single name mapping in the deposited build code.

### 2.2. Query sequences

The type-strain sequences are the 460 records of LactoTypeDB, each classified against a reference that contains it (Table S4, footnote 3). The non-type conspecific sequences were built from the representative type genomes as seeds, and not from a separate accession pull. A family-restricted RefSeq 16S rRNA gene database was searched for the neighbours of each seed, and a neighbour whose assembly returned whole-genome average nucleotide identity of at least 95% to that seed was taken as a conspecific of the seed’s species (Jain et al., 2018). Three of the 431 seeds were quarantined and 107 of the rest returned no search output, leaving 321 searched, of which 215 yielded at least one conspecific. The set is therefore bounded by which species have sequenced non-type relatives rather than by a sampling decision. Those 215 type-strain names fold to 206 species and carry 10,329 sequences of at least 1,200 nt, annotated in 4,898 assemblies.

### 2.3. The three identity constants

The 95% assignment bound above is the lower end of the 95 to 96% range in current use (Chun et al., 2018; Richter and Rosselló-Móra, 2009). Richter and Rosselló-Móra (2009) recommend that whole range from whole-genome comparison and set its upper end at 96%. That lower end is retained here because it is where Goris et al. (2007) put the equivalent of the 70% DNA-DNA hybridization cut-off, and because Konstantinidis and Tiedje (2005) put the same equivalence lower still, at 94% over shared genes. Taking the lower end admits an assembly in the disputed band, so no classifier is credited with declining a query that might have been novel. The 86.5% and 98.65% boundaries cited above decide nothing but the contamination quarantine and the operon concordance report respectively.

### 2.4. Region extraction

Ten regions were scored, the full-length gene and nine shorter ones excised in silico with the trimragged utility of the Unassigner package (Tanes et al., 2024), which locates the forward primer and the reverse-complemented reverse primer and extracts the sequence between them. Table S5 gives every primer with its sequence, *Escherichia coli* coordinates and source (Edgar, 2018b; Herlemann et al., 2011; Sun et al., 2013; Tanes et al., 2024). Klindworth et al. (2013) evaluated the V3-V4 pair and recommended it for community surveys, but designed no primers and credit both of its sequences to Herlemann et al. (2011).

### 2.5. Classification

Unassigner v1.1.1 (Tanes et al., 2024) was run at each of the two thresholds its documentation names as defaults, the hard default (`min_id` 0.975, constant-mismatch-rate algorithm) and the soft default (0.991), a species counting as compatible where its rule-out probability falls below 0.5. No mismatch-position database was supplied, so the variable-mismatch-rate estimator that the released code always instantiates reduces to the constant-mismatch-rate algorithm, its log rate ratio being zero. The search returns every reference sequence above the 90% identity floor, v1.1.1 setting the vsearch `maxaccepts` parameter to 0. The supplemental methods of Tanes et al. set the maximum number of results per query to 5 (Tanes et al., 2024), which the released code does not apply, so no compatible set reported here is truncated. The reference was supplied through the software’s `--type_strain_fasta` argument, which is the only path-based way to pass a curated reference in v1.1.1. The `--db_dir` argument names a managed cache rather than a curated directory, writing the downloaded Living Tree Project release into that directory and overwriting a reference already placed there. LactoTypeDB carries one record per type strain rather than the one record per species that input format expects, so a species with a type subspecies contributes more than one record, a mismatch the fold to parent species under Section 2.6 reconciles.

`assignSpecies` of dada2 v1.38.0 (Callahan, 2025), the release of the package of Callahan et al. (2016) used here, was run with `allowMultiple=TRUE` and `tryRC=FALSE`. `allowMultiple` is the sole non-default argument and it is required, since the default returns NA both where two or more species match and where none does, and those two outcomes are scored separately. Parente et al. (2023) ran `addSpecies` and this work `assignSpecies`, both the exact-match species assignment of the same package.

### 2.6. Scoring

Each compatible set S was scored against the expected species e as resolved (S = {e}), misassigned (one species other than e), ambiguous (two or more) or no call (S empty), the four asserted to sum to the queries scored in each cell. Infraspecific labels were folded to the parent species before scoring. `dada2::assignSpecies` rejects any sequence containing a non-ACGT character, so its denominators are lower than 460 and vary by region (Table S4, footnote 1).

### 2.7. Genus and family assignment

Genus and family were scored separately from species and never stacked with the species outcomes. Both come from `assignTaxonomy`, the naive Bayesian classifier of Wang et al. (2007), run once per region under seed 100 with `minBoot=50`, `tryRC=TRUE` and `outputBootstraps=TRUE`, against SILVA v138.2 (Quast et al., 2013) and against v138.1 for the release comparison. `minBoot=50` is the loosest reporting floor and the setting Parente et al. (2023) ran. It leaves the per-rank bootstrap values intact, so Fig. S1 draws both of its cutoffs, 50 and 80, by filtering those values from that one run rather than from a second run. Only the reference file differs between the two releases. Both were scored by the same scoring function, imported by each run rather than re-implemented for it, over the 434 distinct species and never per genus, against the current roster genus and against the genus SILVA itself records.

### 2.8. Out-of-family probe and V4 groups

One thousand type-strain records from outside the family were sampled with seed 100 from the 19,386 non-*Lactobacillaceae* records of LTP 08_2023. Each was classified by `assignTaxonomy` against SILVA v138.2, scoring whether it was filed in the *Lactobacillaceae*, and by Unassigner at its hard default against LactoTypeDB, with the non-type conspecific sequences as the within-family control reported in Section 3.3. Both ran at full length and in V4, 999 scoring in V4. The V4 groups are the connected components of the reciprocal rule-out graph over the 460 records, defined in Table S6.

### 2.9. Worked example on a published food amplicon survey

The workflow of Table S7 was run end to end on one published food amplicon study, unmodified and by the deposited scripts. The study is Poirier et al. (2023), a longitudinal survey of two fresh meat production lines in France, poultry and raw pork sausage, contributing 435 samples amplified over V3-V4 and sequenced on the Illumina MiSeq platform, held in FoodMicrobionet as study ST283 and in the Sequence Read Archive under accession SRP186244. That study’s curated variant table was taken from the FoodMicrobionet 5.1.2 mindata release (Parente, 2026), a later iteration of the database described by Parente et al. (2022). FoodMicrobionet records the assignment method per study, and for ST283 it is `dada2::assignTaxonomy` against SILVA v138.2. The rows FoodMicrobionet files in the *Lactobacillaceae* were extracted. Sequences carrying the `justConcatenate` run of ambiguous bases were removed, leaving 11,612 distinct sequences of 420 nt, each of them unambiguous ACGT. One record was written per distinct sequence under a content hash, so a variant occurring in several samples is classified once.

Step 1 ran `dada2::assignTaxonomy` against SILVA v138.2 with `minBoot=50` and `tryRC=TRUE`, and the family verdict was resolved at the deepest rank the bootstrap assigned, a query departing the *Lactobacillaceae* lineage at any assigned rank leaving the workflow there. Step 2 ran Unassigner v1.1.1 at its hard default against LactoTypeDB, under the settings of Section 2.5. Step 3 screened the queries step 2 ruled out of every species through four filters in order, being de novo chimera detection with vsearch v2.31.0 (Rognes et al., 2016), a search against the 1,179 16S rRNA gene operons of the representative type genomes, a search against the 10,329 non-type conspecific sequences, and a search against the per-region fallback inserts of Table 1b. Survivors were clustered de novo at the 98.7% species boundary of Yarza et al. (2014), which groups the variants no reference can name into species-level units rather than discarding them, the 3,715 survivors forming 1,549 clusters. Every count reported below is per variant rather than per cluster, so clustering partitions the undescribed fraction without changing any of those counts. Step 4 applied the reporting rule of Table S7 to every query step 1 filed in the family, reading the compatible set from step 2 where there was one and the cascade fate from step 3 where there was none. Where the compatible set was empty, the reported rank follows identity to the nearest described type strain against the published 16S rRNA gene rank boundaries: at or above the 94.5% genus boundary the query is a candidate novel species, between that and the 86.5% family boundary a candidate novel genus, and below the family boundary an exclusion. That identity is step 3’s own search of each query against the reference’s type-strain sequences at this region, rather than the search against the type-genome operons or a separate calculation. Fig. 2d plots the same value, and the deposited scripts assert the two agree for every query and stop on any disagreement.

The input is conditioned on FoodMicrobionet’s own family label, so a variant this study holds that FoodMicrobionet filed elsewhere is absent before step 1 runs. The conditioning removes variants and admits none, thus every count reported from this study is a lower bound. FoodMicrobionet assigns with the same function and the same SILVA release step 1 uses, which bounds the difference between the two family calls to their bootstrap and orientation settings.

### 2.10. Reproducibility

Tool versions are pinned in one place and checked where each tool is called (Table S8).

## 3. RESULTS AND DISCUSSION

### 3.1. LactoTypeDB construction

Every validly published name in the family was taken from LPSN (Parte et al., 2020) and matched against NCBI Taxonomy identifier 33958 (Schoch et al., 2020) on the identifier columns, giving 478 correct names that resolve to 467 type strains across 441 species and 37 genera. The two sources cannot be joined on names, because 19 epithets recur across genera here. For example, *Leuconostoc kimchii* and *Dellaglioa kimchii* are different species that happen to share an epithet. A further 339 LPSN names are synonyms, which the reference does not carry.

Of the 829 candidate assemblies retrieved (Section 2.1), 600 passed the CheckM completeness and contamination thresholds the assembly record reports (Parks et al., 2015) and 99 failed them. A further 130 carry no CheckM values and were recorded as unknown rather than as failures (Table S9a). An assembly deposited under a type strain’s name need not be that strain’s genome, so within-species average nucleotide identity was computed with fastANI v1.34 for the 214 species with more than one candidate (Jain et al., 2018). All 210 species that could be compared were coherent at 97% ANI or above, the lowest observed value being 97.51%.

Seven assemblies that whole-genome contamination scoring passed carried a 16S rRNA gene copy below the 86.5% family-rank boundary of Yarza et al. (2014) while reporting CheckM completeness between 96.68% and 99.35%, four of them at 2.4% contamination or less and three with no contamination value on record. The seven include what were, before quarantine, the top-ranked candidate genomes for *Lactiplantibacillus plantarum* (GCF_051712885.1, lowest copy identity 77.3%) and *Lentilactobacillus buchneri* subsp. *buchneri* (GCF_009495475.1, 74.8%).

Each was quarantined whole rather than by copy (Table 1, footnote 2). Three of the seven were their species’ top-ranked candidate, and two of those species fell back to the next-ranked candidate that passed the screen. *Levilactobacillus tujiorum* was the only species with no other candidate, and lost its genome this way, which is why the genome layer covers 433 species and the 16S rRNA gene layer 434.

16S rRNA gene copies were taken from the PGAP annotation where present (Tatusova et al., 2016) and predicted with barrnap v1.10.5 otherwise (Seemann, 2026). The longest copy passing ribotyper v1.0.5 (Schäffer et al., 2021) was emitted per name, giving 460 sequences across 434 species, of which 454 are distinct (Table 1). Table S2 gives every count with the rule that produced it, including the seven type strains no tier could supply. LPSN also holds an independently deposited 16S rRNA gene accession for 357 of the genome-derived type strains, in close agreement. Across those the median identity is 99.93% and 346 sit at or above the species boundary of Kim et al. (2014), disclosed per record in Table S3. Intragenomic heterogeneity is general in bacteria and reported in *Lactobacillus* (Acinas et al., 2004; Strube, 2021; Větrovský and Baldrian, 2013). Here 71 of the 106 records from a multi-operon genome carry non-identical copies, up to 172 mismatches apart.

### 3.2. What the reference changes

Exact-match species assignment at full length resolved 430 type-strain sequences against LactoTypeDB, misassigned none and left none without a call (Fig. S2). The denominator is 441 of the 460, because `assignSpecies` rejects any sequence carrying a non-ACGT base. Against LTP 08_2023 as published it resolved 44 of the same 441 and returned no call for 397. Only the reference differs between the two.

Parente et al. (2023) resolved 49.9% of this family’s full-length type-strain sequences against a reference they derived from SILVA v138.1. That figure is not comparable with the two above. The reference of Parente et al. (2023) was SILVA v138.1 relabelled rather than assembled from the query sequences, where the sequences scored here are records of LactoTypeDB, and exact matching cannot fail to return a record the reference contains. Self-classification is not equally favourable to all classifiers (Table S4, footnote 3). The held-out measurement is the conspecific column, and the like-for-like comparison is LTP against LactoTypeDB above.

The 11 records left ambiguous at full length are of two kinds. Eight are the four species pairs whose deposited sequences are byte-identical: *Dellaglioa algida* and *D. carnosa*; *Lacticaseibacillus salsurivasis* and *L. suilingensis*; *Levilactobacillus lanxiensis* and *L. lettrarii*; *Secundilactobacillus hailunensis* and *S. silagei*. No method separates those at any length. Three are a shorter deposited record that is an exact substring of a longer record belonging to a sibling species: *Weissella sagaensis* (1,486 nt) within *W. hellenica* (1,576 nt), and *Limosilactobacillus reuteri* subsp. *murium* (1,460 nt) and subsp. *peregrinus* (1,569 nt) within *L. secundus* (1,576 nt). In each of those three the shorter record is contained in the longer one, so exact matching returns both, and the ambiguity is a property of that rule rather than of the marker. Huang et al. (2021) report 100% 16S rRNA gene identity between the type strains of *Lactiplantibacillus plantarum* and *L. argentoratensis*. The two records deposited here are both 1,567 nt and differ at two positions, so that pair separates while the four above do not.

Unassigner downloads the Living Tree Project as its reference, and Tanes et al. (2024) used release LTP 08_2023, as does this work. Against that release as published, Unassigner at its hard default misassigned 98 of 460 type-strain sequences at full length and 1,374 of 10,329 non-type conspecific sequences, the latter being 16S rRNA gene sequences annotated in assemblies other than a species’ representative type genome. With the family’s records replaced by those of LactoTypeDB, no type-strain sequence was misassigned at any region, and misassignment of the conspecific sequences fell to 3 at full length and 31 across all ten (Fig. 1). Single-name calls fell with it, from 215 of 460 to 81 on the type-strain sequences and from 4,293 of 10,329 to 1,698 on the conspecific sequences, the difference moving into compatible sets of two or more. The substitution replaces a single name the marker cannot support with the set of species a query cannot be told apart from. Unassigner assigns more than DADA2 in panel b, which runs in the same direction as Tanes et al. (2024) reported on microbiome reads against a different DADA2 reference (Fig. 3A and Fig. S5 of Tanes et al., 2024, for human body sites and environmental sites respectively), where they attributed much of that difference to type-strain sequences left unannotated at species rank in the reference DADA2 was given. The comparison made here is the within-classifier change from the published release to the substituted one.

**FIG 1.**
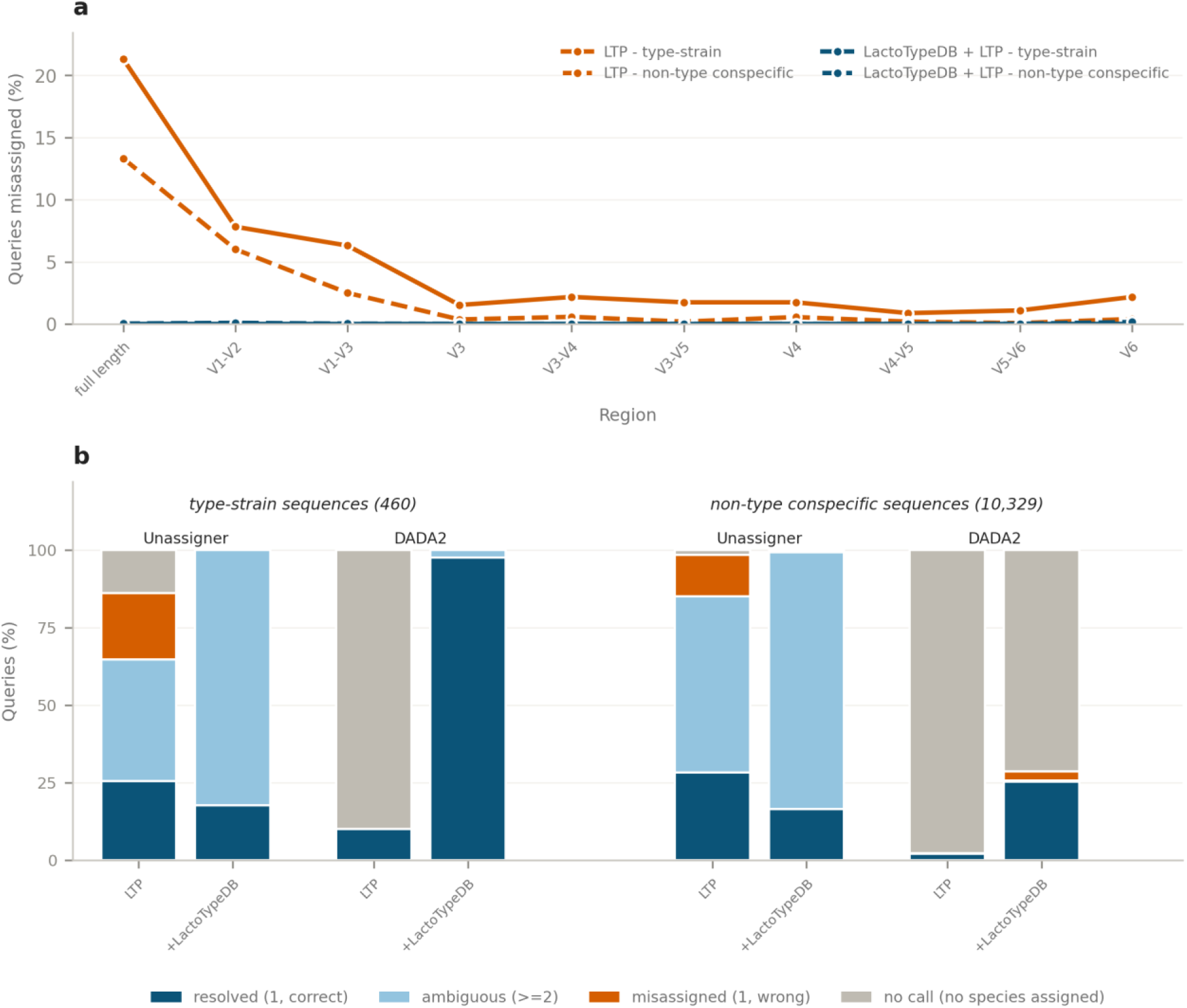
Species-level misassignment against the Living Tree Project reference as published and with the LactoTypeDB records substituted into it. Release LTP 08_2023 as published, shown in the key as “LTP”, against the same release with the family’s records replaced by those of LactoTypeDB, shown as “LactoTypeDB + LTP”. Both sets of query sequences were classified against both references at every region. (a) Percentage of queries misassigned by Unassigner at its hard default (`min_id` 0.975, constant-mismatch-rate algorithm), at full length and nine primer-defined regions, for the type-strain sequences (solid) and the non-type conspecific sequences (dashed). Colour denotes the reference and line style the query sequences. (b) The same substitution for both classifiers at full length, scored as the four mutually exclusive outcomes on both sets of query sequences. Colour denotes the outcome in this panel. LTP 08_2023 contains 376 of the 434 species, so a query whose species is absent cannot match itself and fails for coverage rather than for the content of the record carried. LTP+ 02_2026 holds 32,326 species against 19,509 for LTP 08_2023, each counted as distinct binomials after folding subspecies to the parent (Table S1, footnote 2), and still lacks 44 of the 434, recovering 17 of the 58 that LTP lacked while dropping 3 that LTP held. Panel b compares each classifier against itself across the two references, and the between-classifier difference within it is attributed in Section 3.2. An empty compatible set is an assertion of novelty only for Unassigner, which rules species out. For DADA2 it is the absence of an exact match.

LTP 08_2023 holds 376 of the 434 species, so at most 58 of the 98 full-length misassignments are explained by the query’s species being absent, and at least 40 fell on species the release does hold. Substitution into LTP+ 02_2026 gave the same calls (Table S1).

Filtering against LactoTypeDB left 70 of the 10,329 conspecific sequences with no species-level statement at full length, where exact matching left 7,295 of the 10,228 it accepted. That statement is usually a set rather than a single name, 8,561 of 10,329 at full length and 10,073 in V4. Because the roster carries a record for each of the 434 species that have one, an empty result can no longer mean an unlisted species.

### 3.3. Applying the classifiers in sequence

Unassigner carries no label above species, so it cannot establish family membership, and it returns a set rather than a single name (Tanes et al., 2024). Four steps follow (Table S7). Step 1 classifies one representative sequence per amplicon sequence variant, which is what a DADA2 denoising run emits (Callahan et al., 2016), with `dada2::assignTaxonomy` against SILVA v138.2. It filed all 460 in-family type-strain sequences in the *Lactobacillaceae* at full length and in V4, and none of the 1,000 out-of-family sequences at full length or of the 999 scored in V4. Genus recovery was 423 of the 434 species at full length (Fig. S1), against 416 for the v138.1 release Parente et al. (2023) used, a gap of seven species at each of the four regions they scored. The seven are *Holzapfeliella floricola* and *H. saturejae*, which v138.1 files under *Holzapfelia*, and five *Periweissella* species it files under *Weissella*. Parente et al. (2023) place *Periweissella* with plant and vegetable fermentations and *Weissella* among the generalists, so a survey run on v138.1 reports the two as one genus and cannot separate their distributions.

Across the ten regions v138.2 placed all 460 in the family at full length and at seven of the nine shorter regions, 458 in V5-V6 and 410 in V6, where 8 went to another family and 42 returned no family call. Genus assignment against SILVA needs no revision, other than adjusting for the release version.

Step 2 is `unassign --type_strain_fasta` against LactoTypeDB at the hard default, run on the variants step 1 filed in the family, so LactoTypeDB enters at step 2 alone. A compatible set of one species is a name. Two or more is the set, and in V4 the set is usually one of the 27 groups of Table S6, which lets it be reported by group rather than by an arbitrary member (Nearing et al., 2025). The same out-of-family probe returned an empty compatible set from Unassigner for all 1,000 sequences at full length and all 999 in V4, so without step 1 every read from outside the family reports as a candidate novel *Lactobacillaceae*. The conspecific sequences are the in-family control for that probe, and under the same settings only 70 of them returned an empty set at full length and 35 in V4, so the reference declines an out-of-family query without declining a member of the family. Those 1,000 probe queries are full-length type-strain records rather than amplicon reads, so the complete rejection measured here is the most favourable case available, and a survey working from shorter reads need not reproduce it.

Step 3 takes the queries whose compatible set is empty. Importantly, an empty set against LactoTypeDB is a candidate novel taxon, one of the seven type strains carrying no sequence, or an artefact no reference could match. For the other 434 species it can never be a missing record. A per-region fallback set ships with the release for that second reading. Each of the seven abstaining names carries a sub-full-length 16S rRNA gene locus in its type genome, or an LPSN record the full-length gate rejected. Those sequences were cut to the nine amplicon regions and screened for family coherence by the rule the reference uses. A complete insert was recovered for five of the seven, four of them in V3-V4, and each matches no reference species at its region, so an empty set can be tested before novelty is called. The set is deposited per region beside the reference, and extends species-level screening to those seven for a survey needing the widest coverage the family’s records allow. A survey must still test each empty set against chimeras, further *rrn* copies of a type-strain genome and non-type conspecific sequences before calling any of it novel, then confirm the survivors from genome data, since a new species proposal requires overall genome relatedness index values against the type strains of related species (Chun et al., 2018). Wittouck et al. (2019) labelled eight genome clusters as new species where recovered 16S rRNA gene sequences hit nothing in a type-strain reference restricted to species with no type genome in their data set, and labelled the remaining clusters unidentified where the evidence supported neither call. The type-strain sequences scored here carry no such artefacts by construction, thus step 3 is exercised in Section 3.5 rather than on them.

### 3.4. Where the marker stops, and what novelty then means

In V4, 417 of the 434 species fall into 27 groups the region does not separate, and 258 of those species sit in four groups spanning more than one genus (Table S6). The 27 groups differ in structure: 14 groups are cliques in which every member is inseparable from every other, 11 are chains, which link species pairwise without every member being mutually inseparable, and two are near-cliques. The largest clique holds 22 records resolving to 21 species of *Companilactobacillus*. A chain is not a mutually inseparable set, because a connected component links two species through a third even where those two separate cleanly. The largest chain holds 194 records resolving to 193 species at an edge density of 0.351, so about two thirds of the pairs inside it do separate, and the group names what a query cannot be told apart from rather than a set of equivalent species. That chain includes *Lactobacillus rhamnosus*, the species whose regulatory status the EFSA BIOHAZ Panel revised in 2026 (Section 1), named here under the rule of Section 2.1: even a reference complete to the type strain cannot resolve this case from V4 alone. Tanes et al. (2024) scored 11,917 of 19,791 species as indistinguishable in V4 across all bacteria, at 0 or 1 mismatch rather than by reciprocal rule-out, so that fraction and the one here measure different things.

Resolution tracks which V-regions a region spans rather than its length. With exact matching the V1-V2 region (*Escherichia coli* positions 8 to 338) resolved 412 of 452 type-strain sequences, while the longer V3-V5 region (341 to 926) resolved 287 of 456 (Fig. S3, Table S9). Olivier et al. (2023) nonetheless identified 19 of 20 mock-community species with the full-length 16S rRNA gene, 16 of 20 with V3-V4, and 20 of 20 with the longer 16S-ITS-23S amplicon.

The distinction step 3 rests on is not hypothetical. Parente et al. (2023), surveying this family in foods, found V3-V4 variants that hinted at a different and possibly unknown species of *Holzapfelia* but judged the supporting evidence too scant to act on. That caution was warranted by the reference available to them, and it is the evidence base rather than the judgement that this work changes. Against a reference derived from SILVA that judgement cannot be made, because a variant matching nothing may be novel or may belong to a species the reference does not carry, and the output does not distinguish the two. Against a reference holding a type-anchored record for every species in the family, only the first reading remains, and the variant becomes a candidate to test. Every query scored in Sections 3.2 to 3.4 is a deposited 16S rRNA gene sequence rather than an amplicon read, thus these figures describe the classifiers and the reference rather than the yield of a survey. The next section is that survey.

### 3.5. The workflow run end to end on a food amplicon survey

The workflow of Table S7 was run unmodified on the 11,612 distinct V3-V4 variants that Poirier et al. (2023) recovered from two fresh meat production lines, poultry and raw pork sausage, and that FoodMicrobionet files in the *Lactobacillaceae* (Fig. 2a). Step 1 kept 11,564 of them in the family and released 48, none of which SILVA placed in another family: 38 stopped above genus with the family unresolved and 10 received no placement at all. Step 2 narrowed 213 variants to a single species, returned a bounded set of two or more for 6,740, and ruled out every species for the remaining 4,611. Step 3 withdrew 837 of those 4,611 as further *rrn* copies of a type-strain genome already represented and 59 as matches to a non-type conspecific sequence, and 3,715 survived every screen. De novo chimera detection and the fallback-insert search withdrew none.

**FIG 2.**
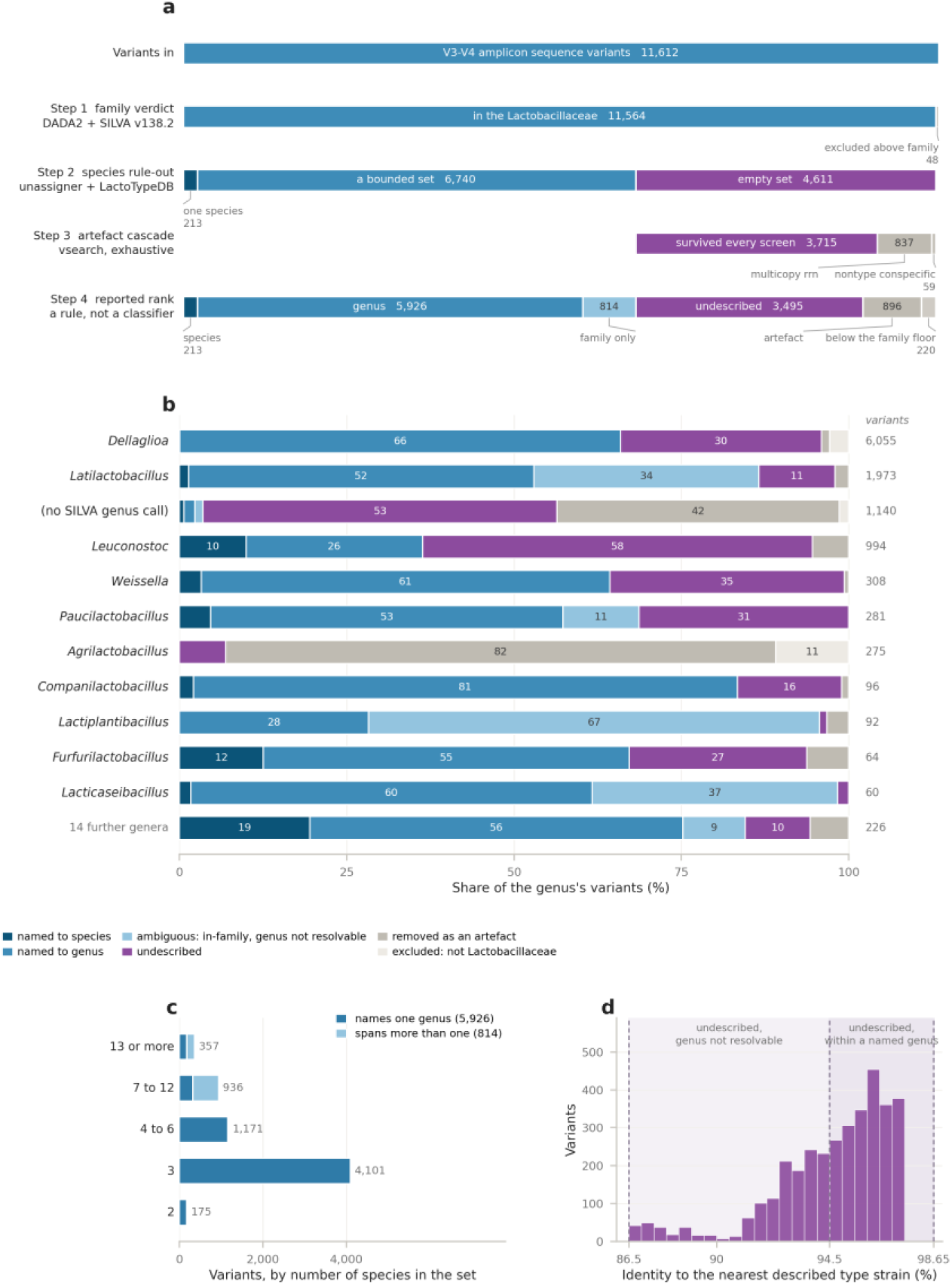
The workflow of Table S7 run end to end on one published food amplicon survey. 11,612 V3-V4 amplicon sequence variants from a longitudinal survey of two fresh meat production lines, poultry and raw pork sausage (Poirier et al., 2023), classified under the workflow unmodified. (a) Every variant’s path through the four steps. Each row partitions what reached it, and the rows are asserted to sum. Step 3 names each class it withdraws. The 59 non-type conspecifics are matches to a described species, withdrawn because the conspecific pool’s 95% average nucleotide identity bound is too permissive to name one (Section 2.3). (b) The rank the workflow reported for each genus SILVA v138.2 assigned at step 1, so each row is the published survey’s own label against the outcome after rule-out. The 11 largest labels are drawn, one of which is the absence of a genus call. The remaining 14 are pooled into one row of 226 variants, being *Fructobacillus*, *Lactobacillus*, *Lapidilactobacillus*, *Lentilactobacillus*, *Levilactobacillus*, *Ligilactobacillus*, *Limosilactobacillus*, *Liquorilactobacillus*, *Loigolactobacillus*, *Paralactobacillus*, *Pediococcus*, *Periweissella*, *Secundilactobacillus* and the uncultured SILVA label HT002. (c) The number of species in the compatible set, for the 6,740 variants the marker narrowed to a set. Colour separates the sets resolving to one genus from those spanning more. (d) Identity to the nearest described type strain, for the 3,495 variants compatible with no described species. Bars are counts in 0.5% bins. Shading gives the two bands of the reporting rule, and the dashed lines mark the 86.5% family, 94.5% genus and 98.65% species boundaries. Median 95.0%, range 86.5 to 97.4%. The identity boundaries derive from the full-length 16S rRNA gene and apply here to a fragment (Table S7, footnote 5). Per-neighbourhood detail is Fig. S4.

Step 4 reported a species for 213 variants, a genus for 5,926, the family alone for 814, and an undescribed candidate for 3,495, with 896 withdrawn as artefacts and 220 excluded below the family floor. The rank reported for a bounded set follows from the set itself. Of the 6,740 sets, 5,926 name one genus and are reported at genus, and the 814 that span more than one are reported at family (Fig. 2c). Every one of the 4,101 sets of exactly three species names one genus, whereas 615 of the 936 sets of seven to 12 span more, thus the reported rank follows what the set contains rather than how large it is.

Of the 3,495 undescribed candidates, 2,111 sit at or above the 94.5% genus boundary and are candidate novel species, and 1,384 fall between that boundary and the 86.5% family boundary and are candidate novel genera (Fig. 2d). Candidate identity to the nearest described type strain has a median of 95.0% and a range of 86.5 to 97.4%. No candidate sits above 97.4%, because a variant closer than that to a described type strain was compatible with it and left step 2 with a name or a set. The 3,495 fall into 22 neighbourhoods by the genus of their nearest described type strain, from 1,434 candidates nearest *Dellaglioa* to a single candidate nearest *Agrilactobacillus* (Fig. S4). The largest neighbourhood sits next to *Dellaglioa algida*, which Poirier et al. (2023) describe as a psychrotrophic meat-borne spoilage organism that becomes dominant during storage, so the undescribed load concentrates around the very species the survey was tracking. In 6 of the 22 neighbourhoods no candidate’s nearest relative resolves to one genus at this region, so those candidates cannot be named even to genus from V3-V4 and are reported as undescribed *Lactobacillaceae*. Splitting the candidates by neighbourhood rather than pooling them is what makes them a target rather than a residue. A laboratory seeking new isolates of industrial interest from this system can return to the samples carrying the *Dellaglioa* neighbourhood rather than to all 435, because the survey’s own sample table records the samples each variant was detected in. Fifty-nine variants matched a non-type conspecific sequence at the rule-out threshold and were withdrawn from the novelty pool. Each is a described species whose 16S rRNA gene differs from that of its type strain, rather than an artefact of sequencing or assembly. The conspecific pool is admitted at 95% whole-genome average nucleotide identity, the permissive end of the 95 to 96% range in current use (Chun et al., 2018; Richter and Rosselló-Móra, 2009), and that end was taken so the false-novelty test would admit assemblies in the disputed band and credit no classifier with declining a query that might have been novel. The bound is therefore strong enough to withdraw a variant from the novelty pool and too permissive to name its species. A species claim on these variants requires digital DNA-DNA hybridization against the matched assemblies (Meier-Kolthoff et al., 2013), which is outside the scope of an amplicon workflow, so they are reported as withdrawn and carried no further. Those 59 matched their conspecific sequence at 97.6 to 98.6% identity over the V3-V4 fragment, which clears the 97.5% rule-out threshold their type strain failed and is what withdraws them, and which names no species on its own. The withdrawal rule holds at the pool’s own bound rather than at the values this study happened to draw.

The genus label the published survey already carries, which is the step 1 SILVA call, is plotted against the rank the workflow reported after rule-out, so each row shows how that genus’s variants were reclassified (Fig. 2b). For *Dellaglioa*, 3,990 of 6,055 variants hold at genus while 1,818 become undescribed candidates and 2 resolve to a species. For *Leuconostoc*, 99 of 994 resolve to a species and 579 become candidates. Where SILVA returned no genus at all, 481 of 1,140 variants proved to be artefacts the cascade withdrew, and for *Agrilactobacillus* 226 of 275 did. Consequently, the artefact load concentrates on the genus labels the marker cannot support. A species name was returned for 213 of the 11,612 variants (1.8%) and a genus for 5,926 (51.0%), against a published survey that reported genus and no more, and the gap between those two figures is the marker’s limit at V3-V4 rather than the reference’s. For a survey admitting isolates to registers such as the Inventory of Microbial Food Cultures, the practical change is that a species call can be trusted where the marker allows it, and a query matching nothing becomes a candidate to test rather than a limitation of the reference to work around.

## 4. CONCLUSIONS

A 16S rRNA gene survey can return only the names that are already in its reference database. Where a species is missing, the result can be a plausible but wrong name that looks no different to a correct assignment. This is due to two separate problems tangling together: how complete the reference is, and how much the marker itself can resolve. LactoTypeDB addresses this for the *Lactobacillaceae* by anchoring each record to a nomenclatural type strain and by regenerating the database from current nomenclature each time it is built, which matters for a family that has been reclassified faster than general-purpose references can keep up with. We substituted it into the Living Tree Project, the standard reference used by established classification software.

Neither the Living Tree Project nor that software is redistributed here. Doing so eliminated misassignment of the family’s type strains at every region tested and cut misassignment of 10,329 other sequences from the same species from 1,374 down to just 3 at full length. This was a reduction of more than a hundredfold. In the widely used V4 region most species cannot be separated, so a survey should report the distinguishable group rather than assign an arbitrary species name. *Lactobacillus rhamnosus*, the species whose regulatory status the EFSA BIOHAZ Panel revised in 2026, illustrates the limit directly: it now carries a type-anchored record in LactoTypeDB, but at V4 it sits in the largest ambiguity group the reference contains, 193 species spanning 16 genera that this region cannot fully separate. A survey using V4 cannot resolve that regulatory case from the marker alone, but it can now say so explicitly rather than silently returning a name. Run on a published survey of two meat production lines the workflow returned a species for 213 of 11,612 distinct sequences and a genus for 5,926 and referred 3,495 for novelty testing rather than naming them. Because every species in the roster now carries a record, a sequence that matches nothing becomes a candidate to test for a new species, not a gap in the reference. Those candidates are a prospecting target rather than a residue, because the workflow reports them by the genus their nearest described relative belongs to. In the meat survey the largest group sat next to *Dellaglioa algida*, the spoilage organism that study was tracking, which gives a laboratory prospecting for industrially useful strains a named target and the samples that carry it. The practical effect is a more scrupulous account of what a fermentation or a spoilage community contains: a survey can report a species where the marker supports one, the group of species where it does not, and an explicit candidate where the query matches nothing, instead of returning a single name in all three cases and leaving a reader unable to tell them apart. Because the build reads only a name roster and the type-strain genomes behind it, the same procedure extends to any other family a food survey needs at species level.

## CRediT authorship contribution statement

**Scott A. Oliphant:** Conceptualization, Data curation, Formal analysis, Investigation, Methodology, Software, Validation, Visualization, Writing – original draft, Writing – review and editing. **Jennifer M. Gardner:** Supervision, Writing – review and editing. **Vladimir Jiranek:** Funding acquisition, Resources, Supervision, Writing – review and editing. **Krista M. Sumby:** Conceptualization, Methodology, Project administration, Supervision, Validation, Writing – review and editing.

## ACKNOWLEDGEMENTS

We thank the curators of the List of Prokaryotic names with Standing in Nomenclature and of the All-Species Living Tree Project, whose maintained resources this work builds upon.

## DATA AVAILABILITY

LactoTypeDB v1.0.0, the code that builds it, the amplicon workflow of Table S7, the code that regenerates every reference that workflow reads, its validation suite and the dated, sha256-pinned snapshot of the LPSN and NCBI inputs are deposited at https://github.com/solipha/lactotypedb. The workflow runs from a clone of that repository against the deposited release by one command, `classify_amplicons.sh`, which takes a query FASTA, the amplicon region those sequences span and an output directory. It resolves every reference it needs from the repository and writes tables: one reported rank per variant, the per-step decisions behind it, and the totals. The region argument selects which per-region fallback set and which collapse groups apply. The release carries the FASTA of 460 sequences, 19 tables and a manifest giving a sha256 for each of those and for every source file, so a rebuild is checkable byte for byte. Every deposited sequence is an existing public record, tabulated in Table S3. The type-strain genome assemblies the build reads are deposited on Zenodo alongside the source archive, because a fresh re-fetch by accession is not guaranteed to be byte-identical: GenBank assemblies are revised in place. The workflow orchestrates dada2 (Callahan, 2025; Callahan et al., 2016), Unassigner (Tanes et al., 2024) and vsearch (Rognes et al., 2016) as external tools and redistributes none of them. The 220-cell scoring table (Table S9) is deposited on Zenodo with the source archive under https://doi.org/10.5281/zenodo.20673517. The worked example’s derived tables are regenerated by running the deposited workflow.

ZENODO_MANIFEST.md records the contents of that deposit. The repository and the Zenodo record are public. The amplicon data reanalysed here are those of Poirier et al. (2023), held in the Sequence Read Archive under study accession SRP186244 (BioProject PRJNA522361) and in FoodMicrobionet as study ST283; the curated variant table was read from the FoodMicrobionet 5.1.2 mindata release (Parente, 2026).

## Declaration of generative AI and AI-assisted technologies

During the preparation of this work, the authors used Claude Code (Anthropic, Claude Opus 4.7) to assist in writing the R, Python and shell code of the LactoTypeDB build pipeline, the amplicon workflow and the analysis and figure scripts. After using this tool, the authors reviewed and edited the content as needed and take full responsibility for the content of the published article.

