## Supplementary Figures for "LactoTypeDB: a regenerable, type-anchored 16S rRNA gene reference for species-level identification of the *Lactobacillaceae* in foods"

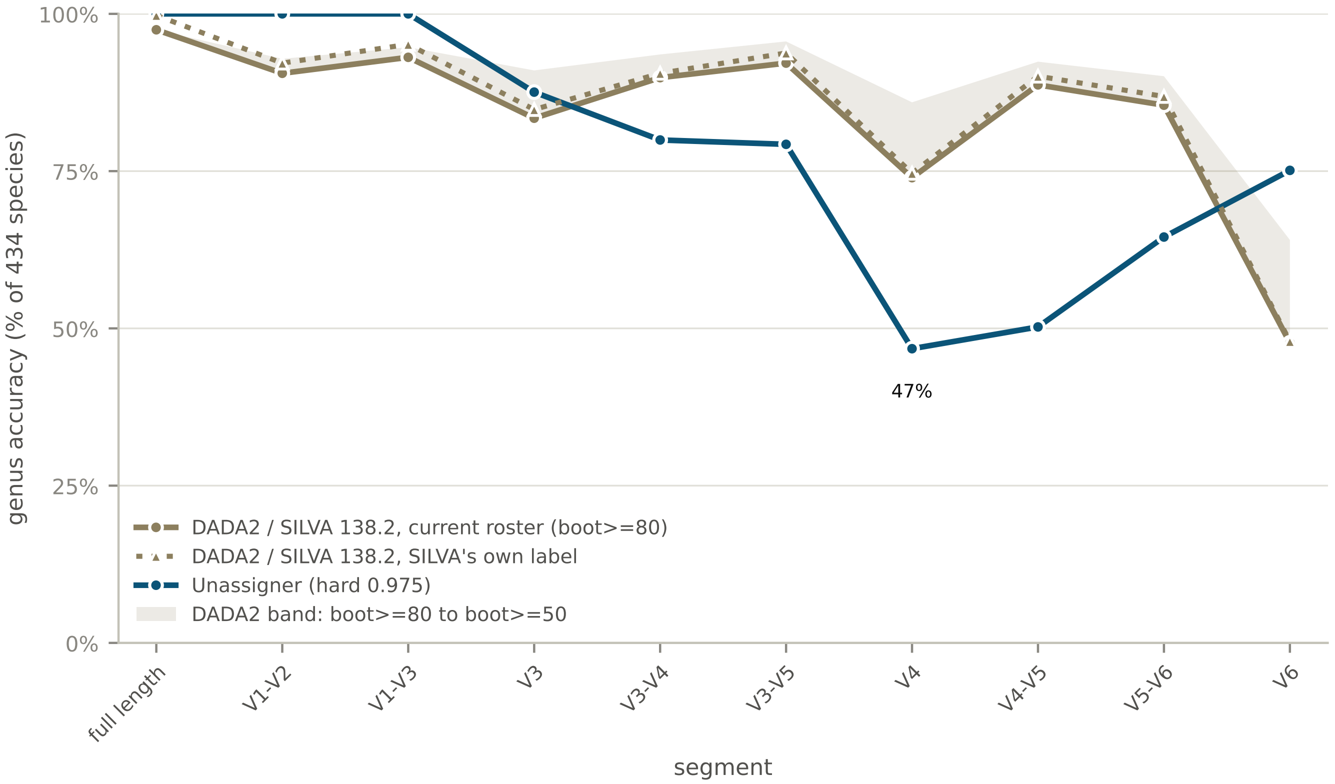


**FIG S1. Genus-level accuracy of two classifiers by region.**

Per-region genus accuracy on the 434 distinct species, infraspecific names folded to the parent. DADA2 calls genus by `assignTaxonomy` against SILVA v138.2 at its documented bootstrap cutoffs, drawn as a solid line at `minBoot` 80 with a shaded band to `minBoot` 50, and as a dotted line where the same calls are scored against the genus SILVA records rather than against the current roster. Unassigner calls genus by folding its hard-default compatible set to genus. A set spanning two genera scores genus-ambiguous.

DADA2 draws on all of SILVA v138.2 and can place a query outside the family. Unassigner folds a compatible set over the 460 LactoTypeDB records, so it draws on the family's 37 genera and its values are conditional on the family membership step 1 establishes.

SILVA v138.2 carries the current genus for 423 of the 434 species and a pre-2020 genus for the other 11, which caps its accuracy against the current roster at every region. The gap between the two DADA2 lines is that nomenclatural lag. At full length DADA2 recovers SILVA's own genus label for 433 of the 434 species. The 11 cover the post-2020 splits SILVA has not adopted, being *Acetilactobacillus*, *Convivina*, *Daquilactobacillus*, *Eupransor*, *Nicoliella*, *Philodulcilactobacillus*, *Xylocopilactobacillus*, and the boundary between *Levilactobacillus* and *Secundilactobacillus*.

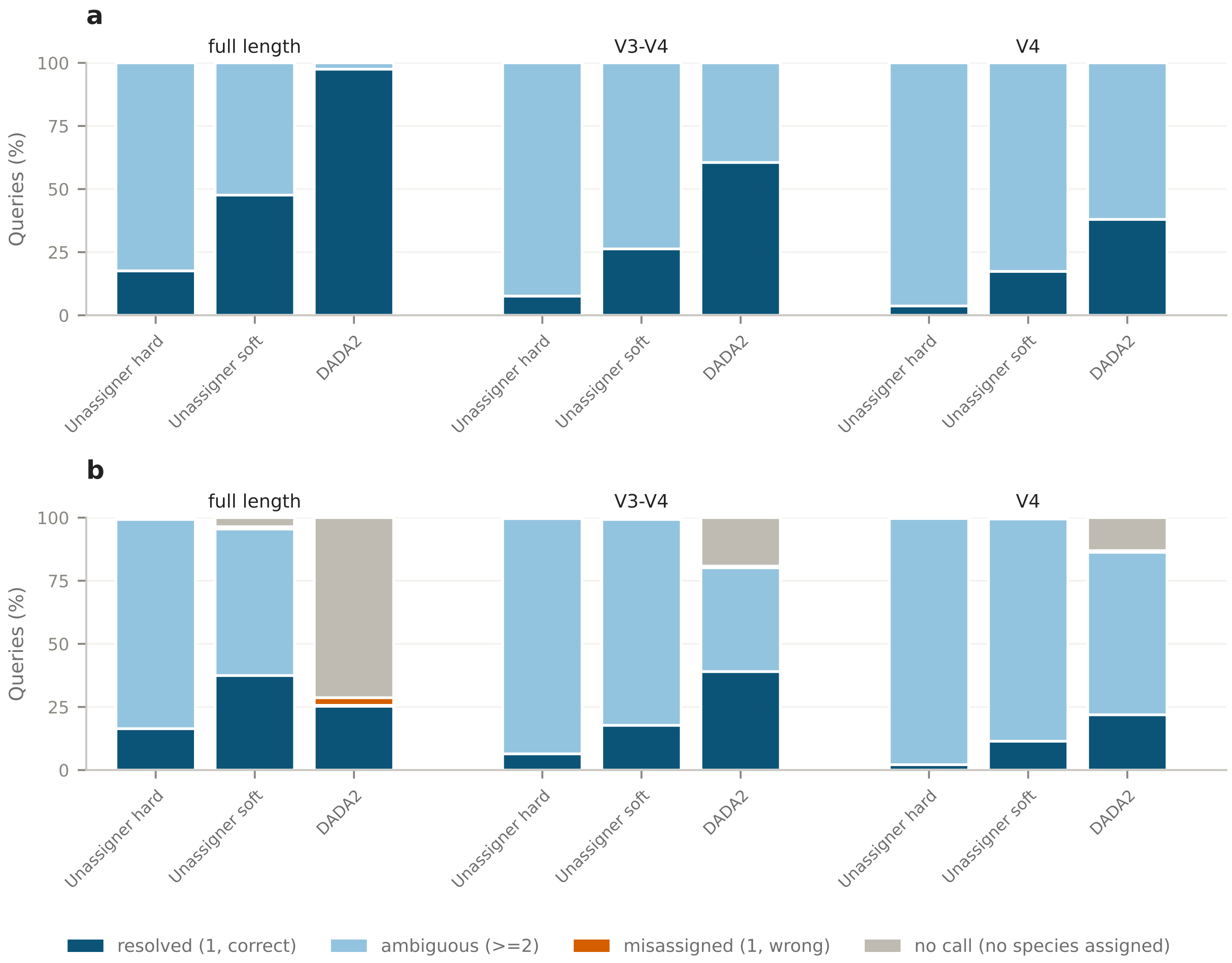


**FIG S2. Four-outcome composition of the species-level call for three classification configurations at three regions.**

Four mutually exclusive outcomes per query, being resolved, ambiguous, misassigned and no call, stacked to 100%, for three configurations against LactoTypeDB, at full length and the two regions most often amplified in community surveys, V3-V4 and V4. (a) the 460 type-strain sequences, self-classified. (b) the 10,329 non-type conspecific sequences, the held-out measurement. All ten regions are Fig. S3.

The species that the V4 region does not separate are given in Table S6, and the four species pairs carrying byte-identical full-length sequences are named in the text.

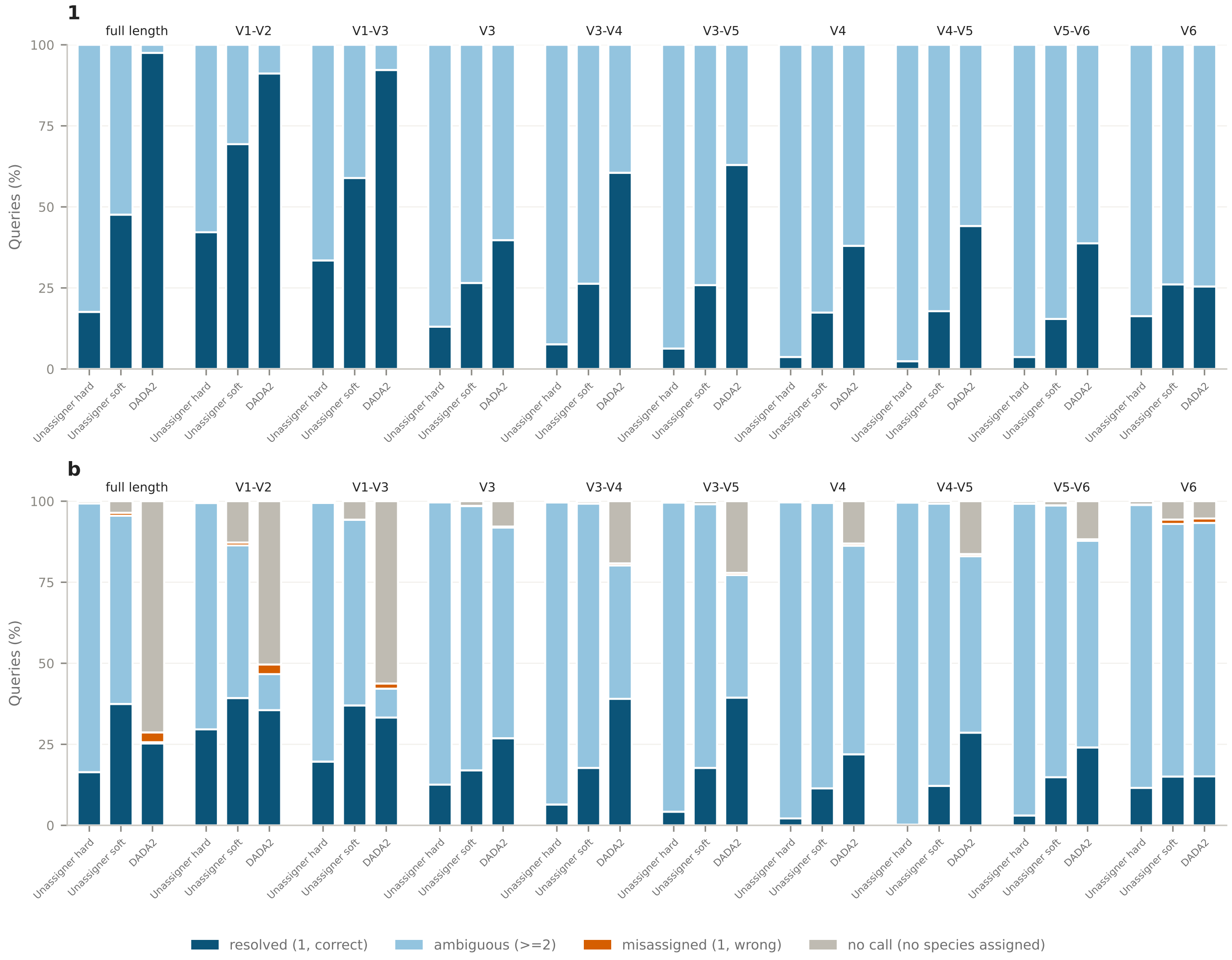


**FIG S3. Four-outcome composition of the species-level call at all ten regions.**

Four mutually exclusive outcomes per query, being resolved, ambiguous, misassigned and no call, stacked to 100%, for three classification configurations against LactoTypeDB: Unassigner at its hard default, Unassigner at its soft default, and `assignSpecies` of DADA2. Both sets of query sequences are shown, the 460 type-strain sequences self-classified and the 10,329 non-type conspecific sequences held out, at the full-length 16S rRNA gene and the nine primer-defined regions of Table S5.

Fig. S2 carries the same three configurations at full length, V3-V4 and V4 alone.

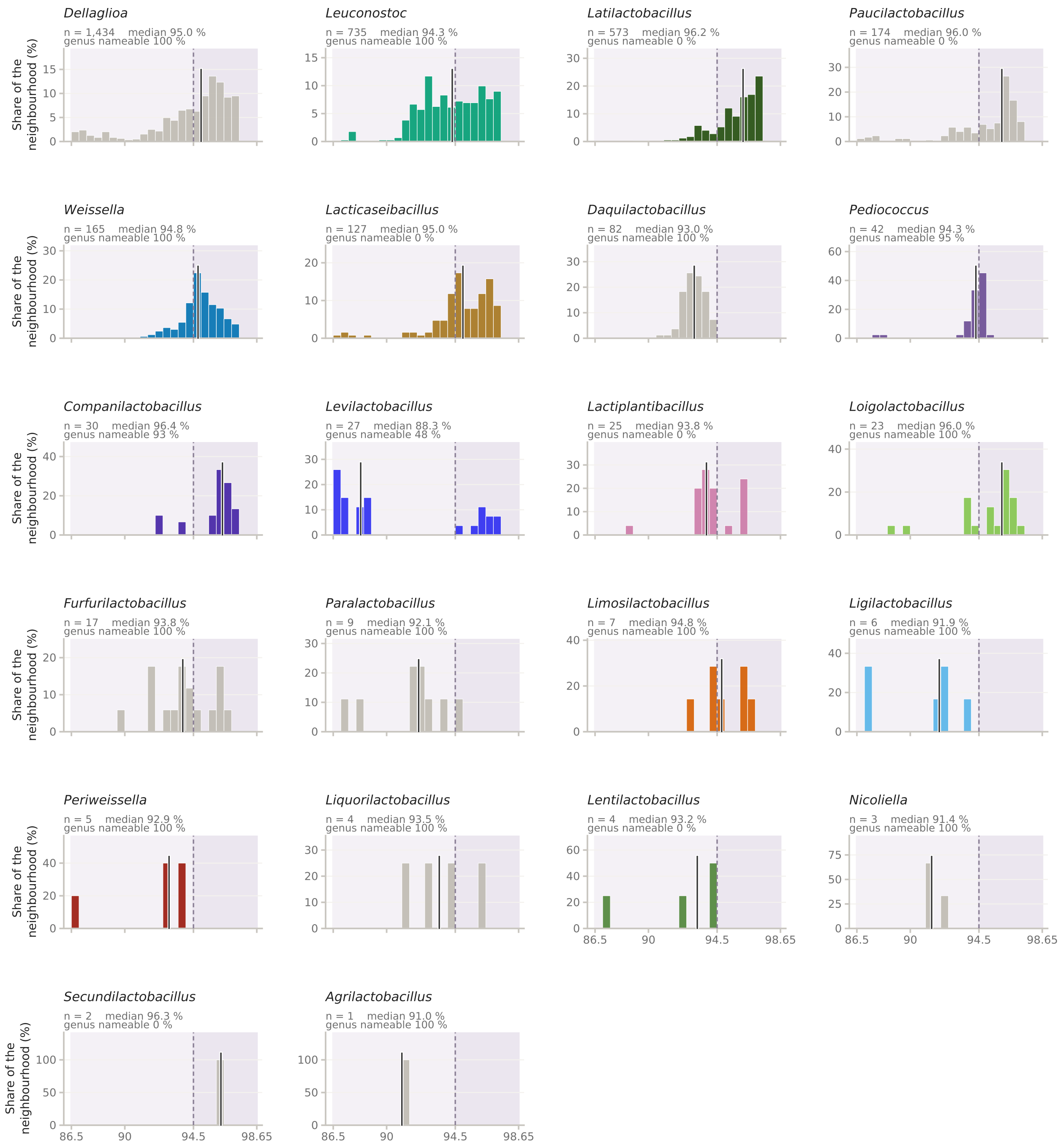


**FIG S4. Every undescribed candidate of Fig. 2, split by the described neighbourhood it sits next to.**

The 3,495 candidates of Fig. 2d, divided among all 22 neighbourhoods, one cell per genus of the nearest described type strain. Bars are the share of each neighbourhood, so shapes compare across cells of very different size, and each heading carries its own count, median and attribution share. Neighbourhoods run from 1 to 1,434 candidates. The vertical rule is the median and the shading gives the two bands of the reporting rule.

Genus nameable is the share of a neighbourhood's candidates whose nearest relative resolves to one genus at this region. A low share marks a neighbourhood the region cannot pin to one genus.
